# Pentoxifylline, a Non-Antibiotic Drug, Promotes Adaptation of Clinical Isolates of *Staphylococcus aureus* via Modulating c-di-AMP Signaling

**DOI:** 10.1101/2025.09.09.675175

**Authors:** Jinru Xie, Lei Wang, Xiukun Wang, Xinyi Yang, Xinxin Hu, Chunjie Xu, Yao Meng, Jianhua Liu, Congran Li, Xuefu You, Guoqing Li

## Abstract

This study investigated how *Staphylococcus aureus* adapts under pentoxifylline (PTX) therapy through analyzing three phylogenetically related isolates (L1, L2, L3) from a bacteremia patient. Whole genome sequencing revealed L1 contained a 2400-bp insertion disrupting *nupC*, while L3 harbored mutations in the anti-Shine-Dalgarno sequences of 16S rRNA. Each isolate evolved distinct mechanisms to reduce c-di-AMP levels, accompanied by changes in survival ability and virulence. Molecular analysis demonstrated PTX noncompetitively inhibits GdpP phosphodiesterase, directly modulating cyclic diadenosine monophosphate (c-di-AMP) signaling. Our findings provide novel insights into how non-antibiotic medications shape bacterial adaptation through second messenger modulation, highlighting the complex trade-offs between stress resistance and fitness in clinical settings.

**Importance:** This study reveals how pentoxifylline, a non-antibiotic drug, drives Staphylococcus aureus adaptation by modulating c-di-AMP signaling. Our findings highlight the unintended consequences of non-antibiotic medications on bacterial evolution, offering new insights for clinical treatment strategies to mitigate resistance development.

**Highlights:**

- PTX noncompetitively inhibits GdpP, modulating bacterial c-di-AMP signaling
- *S. aureus* evolves distinct mechanisms to reduce c-di-AMP under PTX pressure
- Non-antibiotic medications significantly shape bacterial adaptive trajectories

## Introduction

*Staphylococcus aureus*, a highly pathogenic bacterium, causes numerous severe infectious diseases, with bacteremia being one of its most serious manifestations (1). In clinical practice, bacterial pathogens face dual selective pressures, including therapeutic interventions and host immune responses. These pressures drive adaptive evolution that can fundamentally alter their biological characteristics and pathogenic potential (2, 3).

The clinical management of patients with *S. aureus* infections often involves combination therapies. Beyond antibiotic treatment, adjunctive medications used for underlying conditions may inadvertently affect bacterial adaptation (4-9). In addition to drug pressures, the host immune system presents significant challenges to bacterial survival, forcing bacteria to develop immune evasion strategies (10-14). Under these complex pressures, bacteria must adapt to survive, but such adaptation often comes at a biological cost. In this adaptive process, key molecular pathways play crucial roles. However, how bacteria balance adaptation for survival with potential fitness costs under clinical pressure remains poorly understood.

In this study, we investigated three phylogenetically related *S. aureus* strains isolated from a bacteremia patient undergoing PTX combination therapy to better understand how routine medications drive pathogen adaptation in a clinical setting.

## Result

### Basic information, phylogenetic analysis and whole genome alignment of *S. aureus* isolates

Three *S. aureus* isolates (L1, L2, and L3) were obtained from one bacteremia patient undergoing pentoxifylline (PTX) combination therapy. The chronological timeline of strain isolation and therapeutic interventions is illustrated in Figure 1. The patient L, a 63-year-old male was admitted to the hospital on May 23, 2023 after experiencing fever and generalized body pain for three days. Upon admission, blood tests showed a white blood cell count of 25 × 10^9/L, IL-6 of 176.30 pg/mL, and hs-CRP of 180.1 mg/L, indicating that the patient had a severe bacterial infection. Empirical broad-spectrum antibiotic treatment was carried out prior to pathogen identification. Additionally, PTX was administered to improve microcirculation due to the patient’s Grade III hypertension. After 3 days of initial treatment, amoxicillin was replaced with vancomycin upon the isolation of L1 from blood culture. On the 8th day, L2 and L3 were isolated, prompting the addition of moxifloxacin to the treatment regimen. By the 14th day, infection markers had declined, and no further positive blood cultures were obtained until the patient was discharged on June 16, 2023.

**Figure 1.**
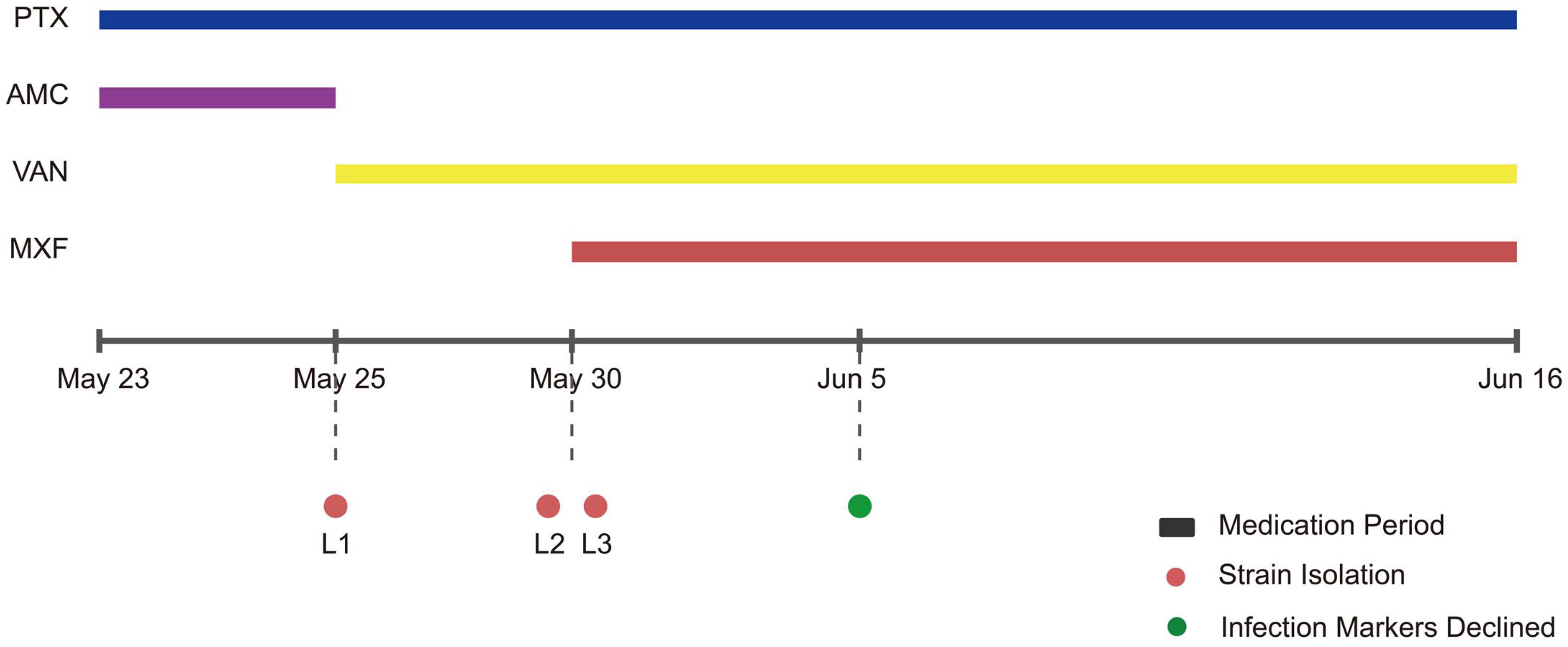
Chronological timeline of therapeutic interventions and clinical events. The timeline illustrates the administration periods of therapeutic agents from May 23 to June 16. Horizontal bars represent treatment durations: PTX (pentoxifylline), AMC (amoxicillin-clavulanate), VAN (vancomycin), and MXF (moxifloxacin). Red circles (L1, L2, L3) indicate timepoints of bacterial strain isolation, while the green circle represents the time at which infection markers declined.

Multilocus sequence typing (MLST) revealed all three isolates belonged to ST965, indicating their substantial genetic similarity. Molecular phylogenetic investigation provided the first insights into the evolutionary relationships among these isolates. The AP distance metrics and NJ method applied to 45 reference *S. aureus* genomes revealed a clustered distribution pattern, suggesting high genomic homology (Figure 2). Branch length calculations illuminated subtle yet significant divergences. L1 displayed a distinctive length of 0.1, whereas both L2 and L3 exhibited shorter lengths, indicating a closer genetic relationship between the latter two strains.

**Figure 2.**
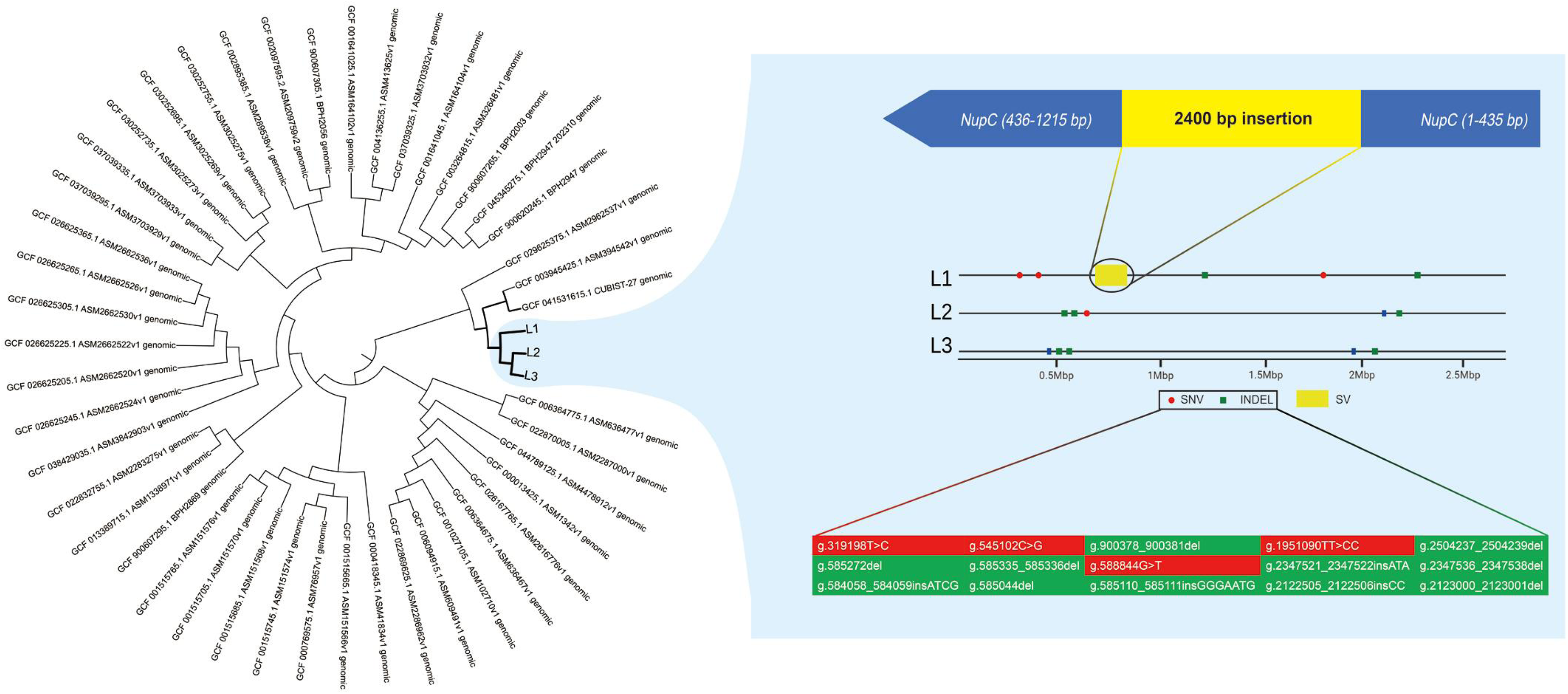
Phylogenetic analysis and genomic comparison of *S. aureus* isolates. Left panel: Neighbor-joining phylogenetic tree based on AP distance metrics incorporating the three patient isolates (L1, L2, L3) and 45 reference *S. aureus* genomes. Right panel: Comparative genomic analysis showing structural variations among the three isolates. The upper diagram illustrates a structural variant (SV) in which a 2400-bp insertion at position 435 bp disrupts *nupC*. Colored blocks in the lower section display detailed sequence variations across the genomes, with SNV marked in red and indels highlighted in green for each isolate (L1, L2, L3).

Comparative whole genome sequence analysis unveiled the molecular basis for the evolutionary divergences (Figure 2). L1 harbored a 2400-bp insertion derived from the *clpC* operon that disrupted *nupC* which encodes a critical mediator of exogenous dNMP transport in nucleoside salvage synthesis (15). In addition to disrupting *nupC* in L1, the 2400-bp insertion introduces extra copies of stress response genes. This genomic modification enhances oxidative stress resistance and alters virulence traits, as previously reported (16-18). In L2, only five single-nucleotide variants (SNVs)/indels were identified, all located in non-coding regions and unlikely to affect physiological functions. L3 carried five SNVs in 16S rRNA coding regions, four of which were situated in the anti-Shine-Dalgarno (anti-SD) sequence, suggesting potential impacts on protein translation (19).

### PTX treatment alters intracellular c-di-AMP concentration in *S. aureus* isolates

As a phosphodiesterase inhibitor, PTX may be related to GdpP, which is a phosphodiesterase that hydrolyzes the second messenger cyclic diadenosine monophosphate (c-di-AMP) to 5’-pApA (20). This enzymatic activity is crucial for maintaining appropriate intracellular c-di-AMP levels in *S. aureus* (21). Molecular docking studies revealed specific interactions between PTX and the DHH domain, which is the catalytic region of GdpP (Figure 3A). PTX demonstrated optimal binding with an energy of -5.9 kcal/mol, indicating stable molecular interactions. Arg499 was identified as a critical stabilizing residue in the GdpP binding pocket, located adjacent to the catalytic metal-coordinating residue Asp497 (22). Subsequently, Isothermal Titration Calorimetry (ITC) was conducted to analyze the interaction between PTX and the DHH-DHHA1 domain of GdpP (GdpP313–656, hereafter referred to as GdpP-C). The titration result revealed a molecular affinity of 33.2 μM between PTX and GdpP-C, comparable to the reported IC_50_ of PTX for phosphodiesterase 3 (PDE3). This suggests PTX has a moderate inhibitory effect on GdpP. This inhibitory effect was further confirmed through enzyme kinetic assays of GdpP-C and its substrate, c-di-AMP, measured both with and without PTX (Figure 3B). Kinetic analysis revealed GdpP-C exhibited similar K_m_ values (101.5 μM vs 101.3 μM) with and without 10 μM PTX, while V_max_ decreased from 48.84 nM/s to 42.60 nM/s. These results demonstrate PTX acts as a noncompetitive inhibitor of GdpP (K_i_ = 68.3 μM).

**Figure 3.**
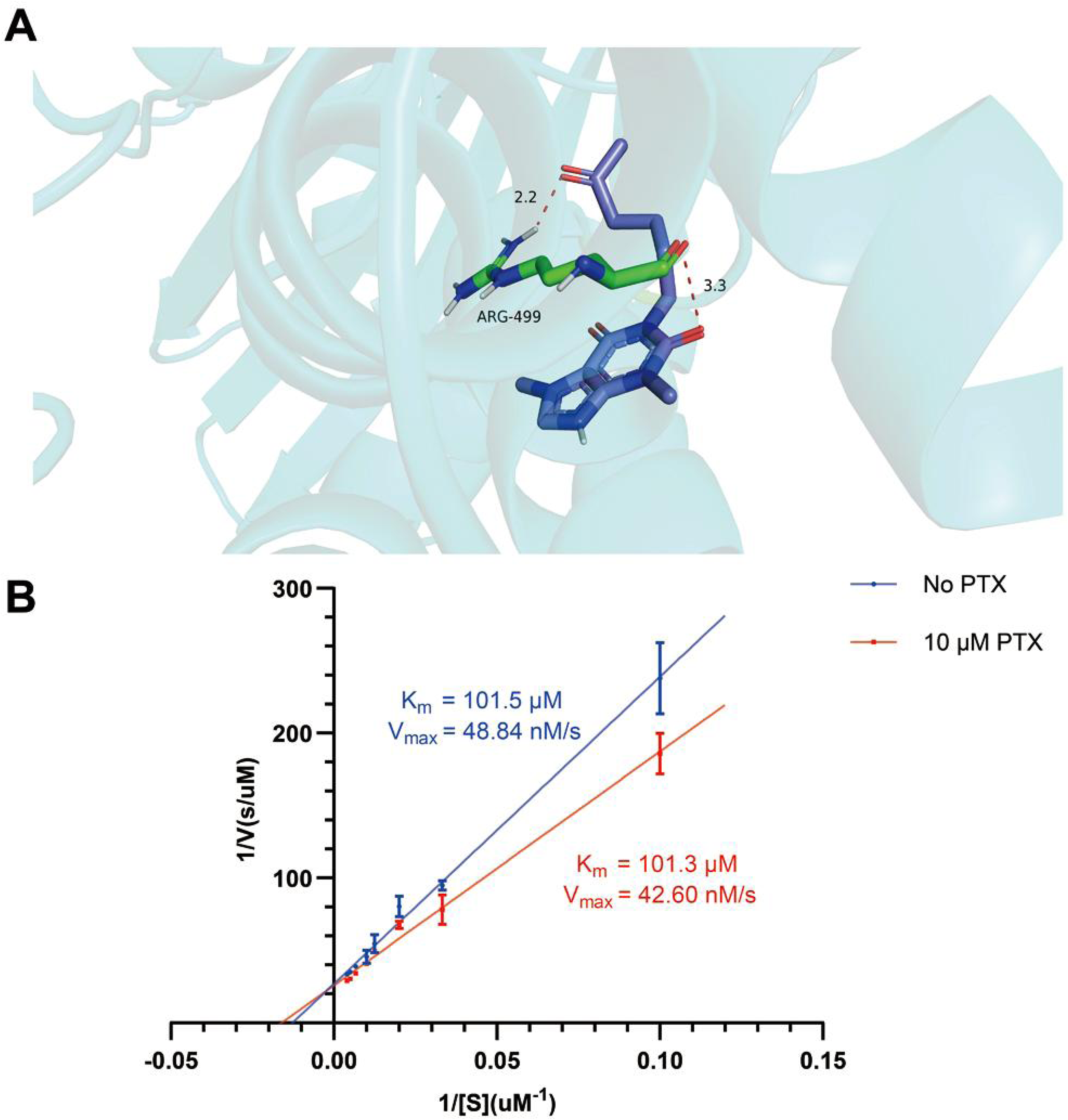
Molecular interaction between PTX and GdpP, and enzymatic kinetics analysis. **(A)** The molecular docking simulation illustrates the binding of PTX (represented by a blue stick model) within the DHH domain of GdpP. ARG-499 forms key stabilizing interactions with PTX, with hydrogen bonding distances of 2.2 Å and 3.3 Å, indicated by dashed lines. The protein backbone is depicted as a cyan ribbon. **(B)** Enzyme kinetic analysis of GdpP-C activity with c-di-AMP as substrate. The reaction velocity (*v*) versus substrate concentration *[S]* curves demonstrate noncompetitive inhibition in the presence of 10 μM PTX (blue) compared to control conditions without PTX (red). Three biological and three technical replicates were performed for each strain. Data are presented as mean ± SD.

Changes in c-di-AMP concentrations in cells after 1 mM PTX exposure at 2, 4, 8, 12, 24, and 48 hours were measured. The results confirmed PTX modulates cellular c-di-AMP levels through GdpP binding (Figure 4A). In all strains, c-di-AMP levels peaked 4 hours post PTX treatment, with L1 showing a 1.79-fold increase, L2 a 4.08-fold increase, and L3 a 1.64-fold increase. Notably, even in the absence of PTX (0-hour time point), L1 and L3 maintained lower c-di-AMP levels compared to L2. Growth analysis under PTX exposure revealed that downregulation of c-di-AMP favored strain growth under PTX pressure (Figure 5B). During the initial 24 hours of 1 mM PTX exposure, bacterial growth exhibited a hierarchical pattern (L1 > L3 > L2). This growth disparity suggests that the reduced c-di-AMP levels in L1 and L3 provided a growth advantage under PTX pressure compared to strains with normal c-di-AMP levels.

**Figure 4.**
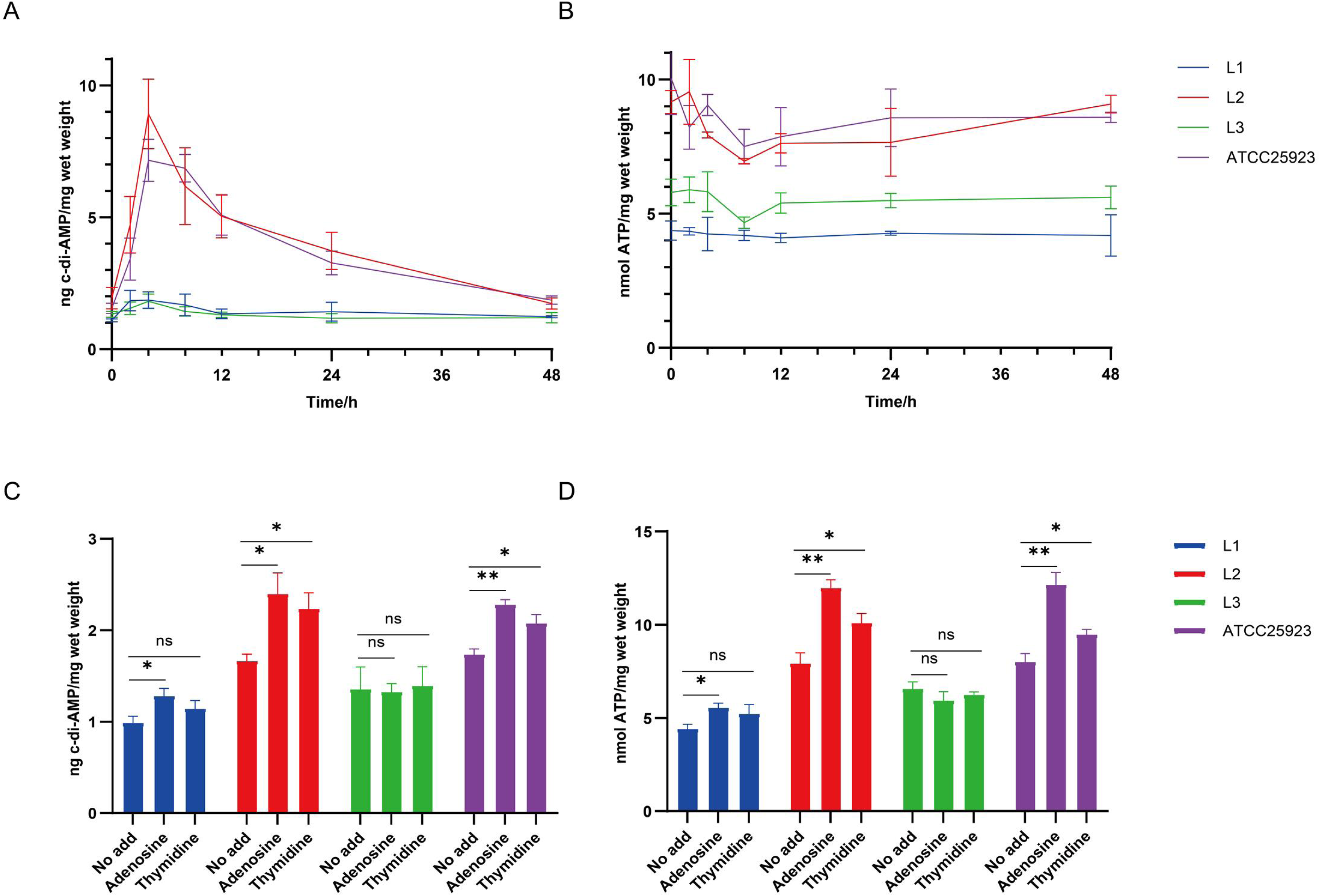
Dynamics of c-di-AMP and ATP levels in *S. aureus* isolates under PTX treatment and nucleoside supplementation. **(A)** Temporal profile of intracellular c-di-AMP concentrations over 48 hours following PTX exposure (1 mM) across isolates L1 (blue), L2 (red), L3 (green), and reference strain ATCC 25923 (purple). Three biological and three technical replicates were performed for each strain. Data are presented as mean ± SD. **(B)** Temporal profile of intracellular ATP concentrations over 48 hours following PTX exposure (1 mM) across isolates L1 (blue), L2 (red), L3 (green), and reference strain ATCC 25923 (purple). Three biological and three technical replicates were performed for each strain. Data are presented as mean ± SD. **(C)** Changes in intracellular c-di-AMP content in isolates L1 (blue), L2 (red), L3 (green), and reference strain ATCC 25923 (purple) after 12 h treatment with different nucleoside supplementations (no addition, 100 μM adenosine, or 100 μM thymidine). *P< 0.05, **P< 0.01, **P< 0.001, by two-tailed t-test. Three biological and three technical replicates were performed for each strain. Data are presented as mean ± SD. **(D)** Changes in intracellular ATP content in isolates L1 (blue), L2 (red), L3 (green), and reference strain ATCC 25923 (purple) after 12 h treatment with different nucleoside supplementations (no addition, 100 μM adenosine, or 100 μM thymidine). *P< 0.05, **P< 0.01, **P< 0.001, by two-tailed t-test. Three biological and three technical replicates were performed for each strain. Data are presented as mean ± SD.

**Figure 5.**
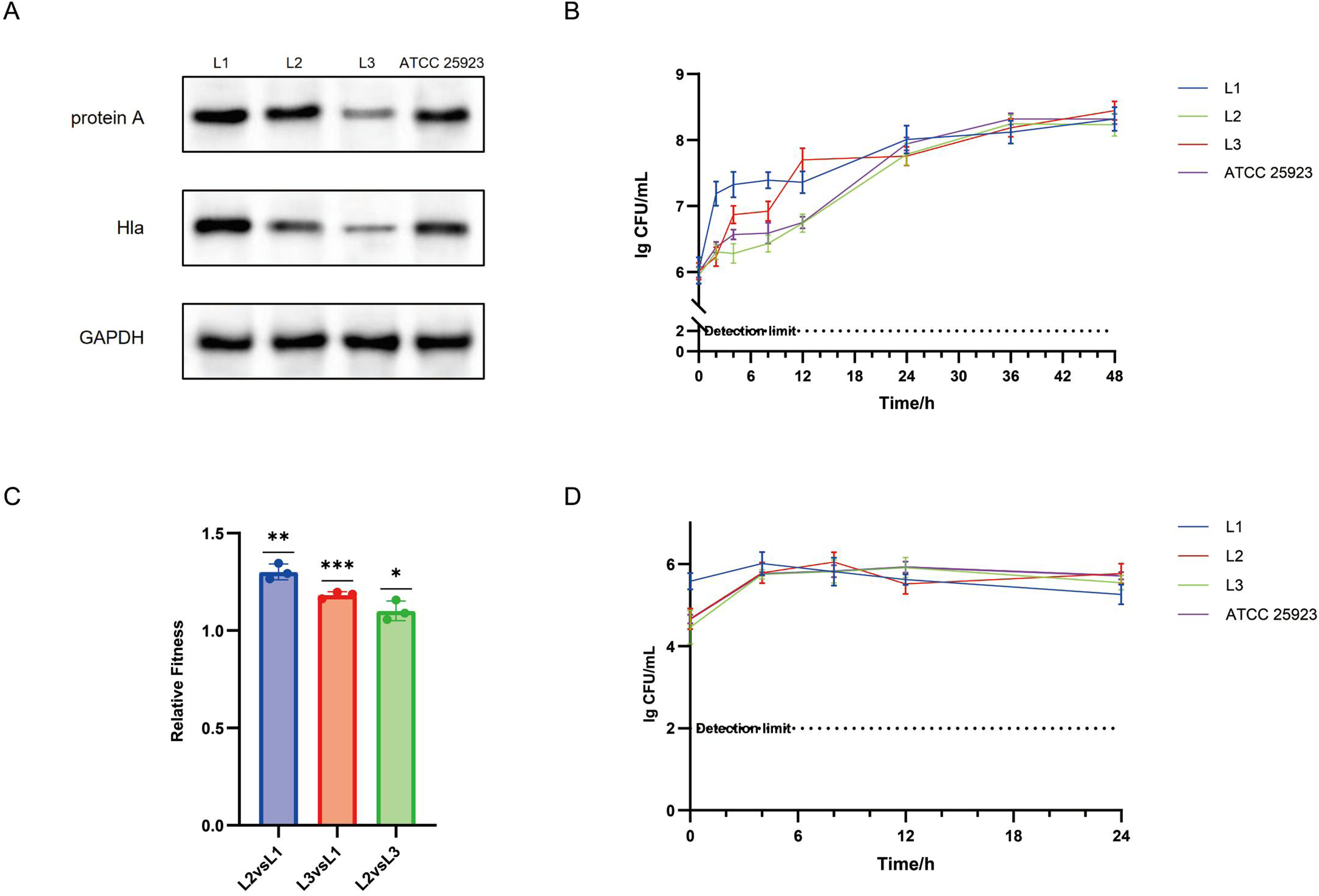
Molecular phenotypic and fitness profiling of PTX-adapted *S. aureus* isolates. **(A)** Protein expression levels of Protein A (SpA), α-hemolysin (Hla), and the loading control GAPDH in L1, L2, L3 and ATCC 25923. **(B**) Growth curves were measured over 48 hours following PTX exposure (1 mM) across isolates L1 (blue), L2 (red), L3 (green), and reference strain ATCC 25923 (purple). Three biological and three technical replicates were performed for each strain. Data are presented as mean ± SD. The dashed line indicates the detection limit of the assay. **(C)** *In vitro* competition assay of isolates L1 (blue), L2 (red) and L3(green). Three biological and six technical replicates were performed for each strain. Data are presented as mean±SD. *P<0.05, ***P<0.0001, by two-tailed t-test. **(D)** The survival curves of isolates L1 (blue), L2 (red), L3 (green), and reference strain ATCC 25923 (purple) in A549 (human-derived lung cancer cells). Three biological and three technical replicates were performed for each strain. Data are presented as mean±SD. The detection limit is indicated by the horizontal dotted line.

### Effects of PTX on intracellular ATP concentration in *S. aureus* isolates

Since ATP serves as the substrate for c-di-AMP synthesis in *S. aureus* (23), we monitored ATP concentration under PTX exposure (Figure 4B). Results showed that ATP level in L1 and L3 remained significantly lower than that in L2. Under PTX exposure, the ATP content of L2 and ATCC25923 was decreased during the initial 8 hours but gradually increased thereafter. This suggests that the low c-di-AMP levels in L1 and L3 may stem from limited substrate availability caused by low ATP concentrations. Conversely, L2, which maintained normal c-di-AMP levels, may experience negative feedback that reduces ATP and stabilizes c-di-AMP level at a lower equilibrium (24). Meanwhile, L1 and L3 maintained consistently low c-di-AMP levels that did not fluctuate significantly.

The low ATP level in L1 may result from the kilobase-scale genomic insertion disrupting *nupC*. Notably, NupC (a nucleoside transporter in the salvage pathway) and ThyA (a thymidylate synthase essential for *de novo* dTMP synthesis) exhibit functional complementarity(25, 26). In L1, *thyA* expression was 5-fold higher than in L2/L3 (Table 1), suggesting compensatory upregulation due to impaired nucleotide metabolism. Disruption of salvage pathway genes (e.g., *nupC*) can reduce ATP levels by both diminishing substrate-driven ATP synthesis (27-29) and increasing ATP consumption (30). Consistent with this, our nucleoside supplementation experiments confirmed altered ATP dynamics in *nupC*-disrupted strains (Figure 4D). Compared to L2, ATP content in L1 did not fluctuate when supplemented with exogenous thymidine, whereas that in L2 showed significant increase. With increased exogenous adenosine, L1 exhibited a slight increase in ATP content, while L2 demonstrated a more pronounced increase. The variations in c-di-AMP content across all three strains were consistent with the changes in ATP levels following the addition of different nucleosides (Figure 4C).

**Table 1.**
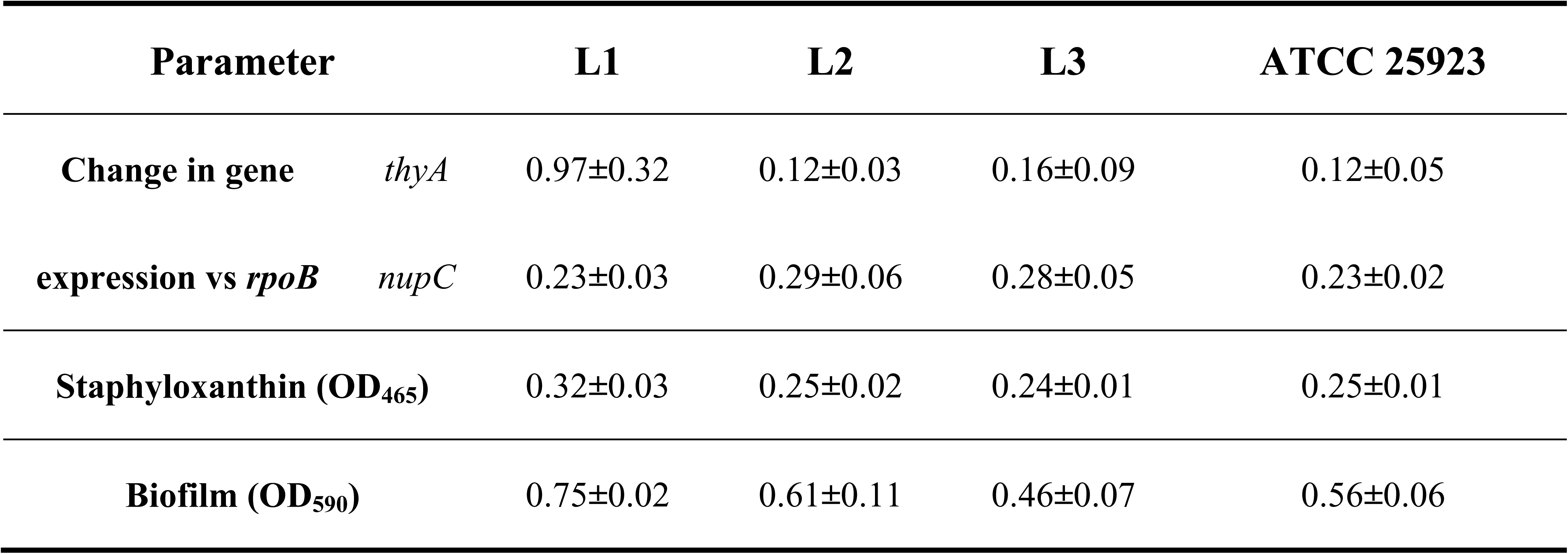
Comparative analysis of nucleoside metabolism-related gene expression and phenotypic characteristics of *S. aureus* isolates.

ATP and c-di-AMP levels in L3 remained unchanged with nucleoside supplementation, suggesting that L3’s mutations limit ATP through mechanisms independent of nucleoside synthesis. Whole-genome alignment revealed that L3’s mutations were concentrated in the anti-Shine Dalgarno (anti-SD) sequence of 16S rRNA. To investigate the functional impact of these mutations, we examined the expression of two SD sequence-dependent proteins in *S. aureus*, Protein A (SpA) and α-hemolysin (Hla). The results showed that L3 exhibited significantly reduced levels of SpA and Hla compared to both L1 and L2 (Figure 5A). This finding indicates that the expression of SD sequence-dependent proteins, including those involved in ATP synthesis, is markedly downregulated in L3, ultimately influencing ATP and c-di-AMP levels.

### PTX-induced adaptation leads to differences in survival capabilities among *S. aureus* isolates

These three isolates displayed varying susceptibility to cotrimoxazole (Trimethoprim/sulfamethoxazole), with minimum inhibitory concentrations (MICs) of <1/19, 4/76, and 2/38 μg/mL for L1, L2, and L3, respectively. All strains were sensitive to other antibiotics, exhibiting identical MIC for oxacillin, vancomycin, linezolid, daptomycin, and clindamycin (data not shown), suggesting that the resistance differences were specific to cotrimoxazole. As previously mentioned, the inhibition of the thymidine salvage pathway likely contributes to it (15, 25, 31, 32). Specifically, when cotrimoxazole suppresses the endogenous thymidine synthesis pathway, which L1 predominantly depends on, it severely compromises bacterial viability (25, 32-34). Competitive fitness assays were performed to evaluate the potential cost of adaptation, revealing notable disparities among the isolates. Both L2 and L3 exhibited significantly greater fitness compared to L1 (Figure 5C). Moreover, when interacting with A549 (pulmonary epithelial cell), L1 demonstrated superior invasion capacity during the initial co-incubation period, while its intracellular survival rates (24-hour time point) remained comparable to the other strains (Figure 5D).

These findings suggest that the distinct adaptive strategies adopted by L1 and L3 under PTX exposure have influenced their survival capability. L1 exhibits stronger invasion ability, while this advantage comes at a substantial cost to its fitness, resulting in a diminished competitive capacity under normal growth conditions.

### *S. aureus* isolates adapted to PTX stress exhibit differential virulence

As mentioned previously, western blot analysis revealed a distinct hierarchical expression pattern of SpA and Hla (L1>L2>L3) (Figure 5A) which are both virulence-related proteins, suggesting that L1 and L2 may possess higher virulence potential than L3. Quantification of staphyloxanthin production by absorbance measurements at 465nm showed that L1 produced a significantly higher concentration of staphyloxanthin compared to L2 and L3 (Table 1). A similar trend was observed in biofilm formation capacity, measured by crystal violet staining at 590nm (Table 1).

*In vivo* studies using a murine infection model elucidated distinct patterns of host adaptation and virulence. Survival analysis revealed that L1- and L2-infected mice exhibited 100% mortality, whereas L3-infected ones maintained a 30% survival rate (Figure 6A). ELISA-based quantification of serum IL-1β revealed significantly higher levels in L2-infected mice at 2 hours post-infection compared to L1- and L3-infected groups (Figure 6B). While both L1 and L2 exhibited peak IL-1β levels at 2 hours post-infection, L3 infection triggered a delayed response, with peak concentrations occurring at 4 hours post-infection. As IL-1β is a major inflammatory mediator (35), its rapid elevation suggests stronger early immune activation in L2-infected hosts. These observations imply that L2’s heightened murine virulence may stem from its ability to provoke excessive inflammatory responses. Quantitative assessment of bacterial burden demonstrated strain-specific tissue tropism, with L1 showing significantly greater pulmonary colonization efficiency. At 72 hours post-infection, L1 achieved bacterial loads approximately two logarithmic units higher than L2 and L3 in the lungs (6.69 vs 4.90 and 4.83 lg CFU/g, respectively) (Figure 6C). This enhanced pulmonary colonization capacity of L1 aligns with its superior virulence observed *in vitro*.

**Figure 6.**
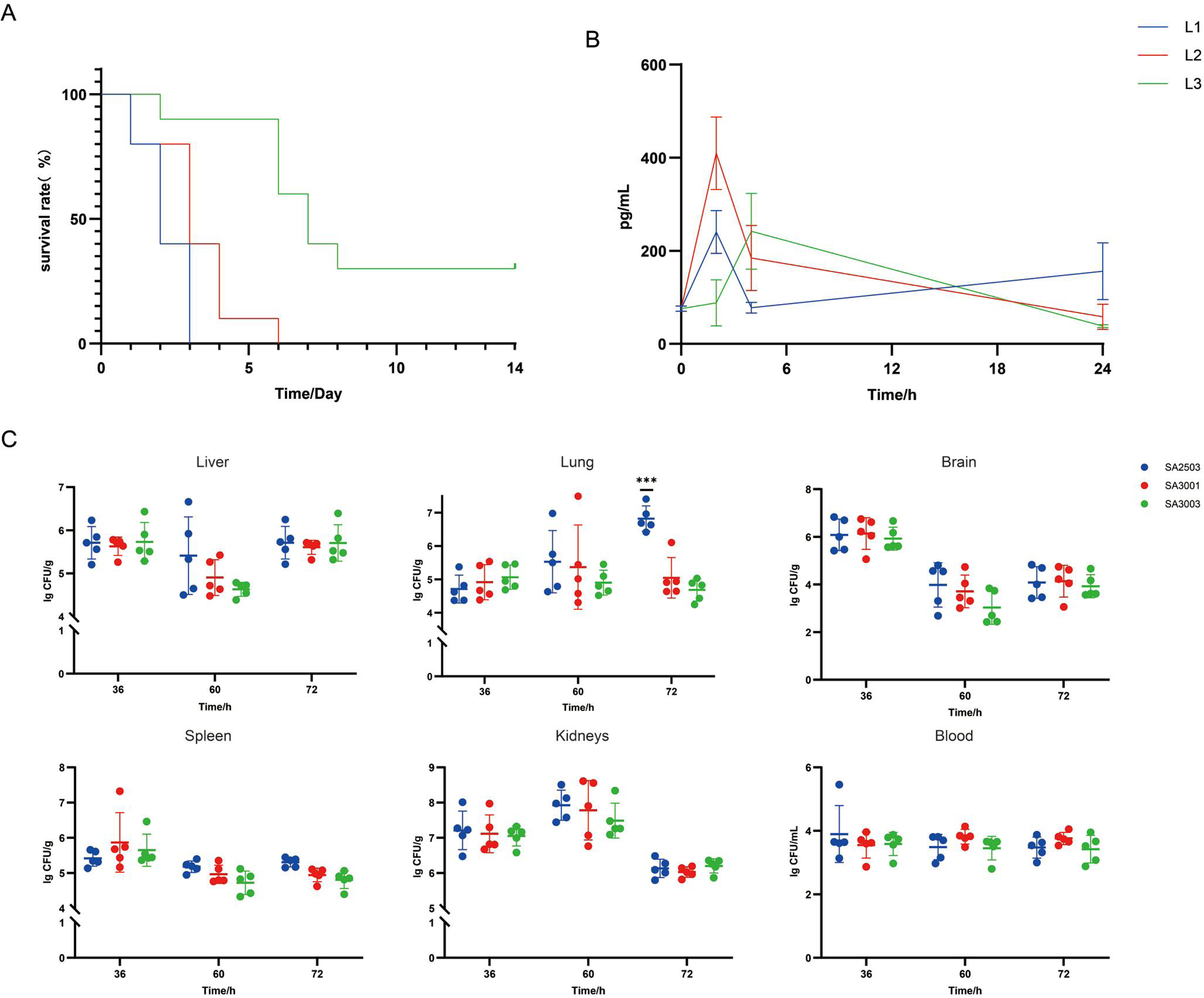
Virulence, inflammatory response, and tissue-specific bacterial burden of the three isolates in a murine infection model. **(A)** Characterization of virulence phenotype of L1 (blue), L2 (red), and L3 (green) over a 14-day observation period using a mouse systemic infection model (n = 10). The intravenous infection dose was approximately 3×10^7^ CFU per mouse. **(B)** Quantification of serum IL-1β concentrations (pg/mL) in mice infected with L1, L2, and L3 over a 24-h period post-infection. The intravenous infection dose was approximately 10^7^ CFU per mouse. Three biological and three technical replicates were performed for each strain. Data are presented as mean ± SD. **(C)** The bacterial loads of L1 (blue), L2 (red), and L3 (green) in the liver, lung, brain, spleen, kidney, and blood of mice, respectively. The intravenous infection dose was approximately 10^7^CFU per mouse. Data are presented as mean ± SD (n = 5). ***P<0.0001, by two-tailed t-test.

In summary, these results indicate that bacterial virulence levels evolved as a result of adaptation to PTX exposure. The observed differences in virulence are primarily driven by the varying c-di-AMP expression levels among the three strains.

## Discussion

This study provides a paradigmatic example of how *Staphylococcus aureus* evolves distinct adaptive strategies under therapeutic and immunological pressures. Through comprehensive analysis of three isolates from a single bacteremia patient undergoing combination therapy, we uncovered diverse adaptive trajectories, offering critical insights into how bacterial pathogens navigate the trade-offs between stress resistance and survival capabilities under clinical pressure.

The growing recognition of non-antibiotic medications’ impact on bacterial physiology and evolution has become pivotal in therapeutic strategies (4-9). Recent studies have demonstrated that commonly prescribed drugs, including antipsychotics, antidepressants, and anti-inflammatory agents, can significantly modulate bacterial gene expression, metabolism, and virulence through non-traditional mechanisms(8, 9, 36). Our study provides the first evidence that pentoxifylline (PTX), a widely used hemorrheologic agent, can shape bacterial adaptive trajectories by directly modulating bacterial c-di-AMP signaling via GdpP inhibition.

The mechanistic basis of L1’s adaptation centers on nucleotide metabolism disruption. Since thymidine transport in bacteria primarily involves NupC, while adenosine transport involves both NupC and NupG, the frameshift mutation of *nupC* in L1 more significantly affects exogenous thymidine transport and related ATP content changes rather than adenosine transport (15). This phenomenon of exogenous nucleoside transporter mutations reducing c-di-AMP levels through disruption of the nucleoside salvage pathway has previously been observed in NupG (37). Here, we identified a novel molecular mechanism linking NupC disruption to c-di-AMP regulation, suggesting more complex interactions between nucleotide metabolism and second messenger signaling than previously recognized.

In contrast, L3 adapted by altering its anti-SD sequence, which decreased SD sequence accessibility and potentially downregulated the expression of proteins dependent on the SD sequence for translation (19). As ATP synthase relies on SD sequence-dependent translation (38), this mutation likely reduced ATP content through the synthase pathway, subsequently lowering c-di-AMP levels. Additionally, c-di-AMP, as a critical second messenger, exerts its physiological functions through interactions with *ydaO*-type riboswitches (39), which are directly connected with SD sequence. The disruption of the anti-SD sequence in L3 may have a feedback effect on c-di-AMP synthesis, further contributing to its downregulation.

In terms of survival adaptability and phenotypes, L1 significantly enhanced virulence at the cost of reduced competitive fitness. Its advantage in the pulmonary environment may be attributed to its enhanced cytotoxicity toward pulmonary cells, as observed in co-incubation experiments *in vitro* (40). Nevertheless, the high energetic demands of maintaining elevated resistance, virulence, and compensatory metabolic pathways likely explain L1’s diminished competitive ability, which ultimately led to its elimination in the second sampling during competition with L2 and L3.

As to L3, it adapted by downregulating protein expression through ribosomal modifications, achieving moderate stress resistance while minimizing energy expenditure. This strategy may also reduce the number of pathogen recognition receptor (PRR) targets (41). Coupled with its low virulence *in vitro*, this likely explains the observed lowest mortality rate *in vivo*.

A particularly intriguing observation was L2. Despite retaining characteristics most similar to the wild-type strain and exhibiting intermediate virulence phenotypes among the three clinical isolates *in vitro*, it triggered the most robust inflammatory responses among the three clinical isolates *in vivo*. This heightened immune activation, potentially mediated by normal c-di-AMP levels compared to the reduced levels in L1 and L3, may explain the comparable mortality rates between L1- and L2-infected animals despite their distinct virulence mechanisms. This finding underscores the complexity of host-pathogen interactions and the potential disconnect between *in vitro* phenotypes and *in vivo* pathogenicity.

The emergence of these diverse adaptive strategies within a single host highlights the remarkable evolutionary plasticity of *S. aureus* and underscores the need for adaptive therapeutic approaches that anticipate bacterial evolution. This is particularly critical in clinical settings where treatment regimens can rapidly shift, creating varying selective pressures. The observed trade-offs between stress resistance and basal growth advantages emphasize the importance of considering evolutionary costs when designing therapeutic strategies.

The rapid emergence of these adaptations raises several critical questions for future investigation. How do bacterial populations maintain diversity in adaptive strategies within a single host? What determines the choice of adaptive trajectory? How can we predict and potentially guide bacterial adaptation in clinical settings? Future studies should focus on developing methods to predict likely adaptive trajectories under specific therapeutic regimens, understanding the role of host factors in shaping bacterial adaptation, and investigating therapeutic strategies that exploit evolutionary trade-offs.

In conclusion, our study provides unprecedented insights into bacterial adaptation under therapeutic pressure, particularly highlighting the complex role of non-antibiotic medications in shaping bacterial evolution. The identification of novel molecular mechanisms related to *S. aureus* c-di-AMP regulation challenges existing paradigms and suggests new considerations for therapeutic strategies. These findings emphasize the need to carefully consider the effects of non-antibiotic medications on bacterial adaptation when designing treatment regimens, potentially opening new avenues for optimizing therapeutic outcomes in infectious diseases.

## MATERIALS AND METHODS

### Bacterial Isolates and Growth Conditions

The *S. aureus* isolates (L1, L2, and L3) used in this study were collected from a 63-year-old male patient at the First Affiliated Hospital, Hebei North University (Hebei, China). Positive blood culture samples were streaked onto blood agar plates, then incubated at 37°C for 24 hours. The *S. aureus* stains isolated on May 25, 2023 (named L1) and May 30, 2023 (named L2 and L3) were recovered by streaking on Luria-Bertani (LB) agar plates, and identified by VITEK®2 Compact System (bioMérieux, France) and Autof MS1000 automated mass spectrometry microbial identification system (Autubio, China). Single colonies of each sample were randomly picked and stored at −80°C in the Chinese Academy of Medical Sciences Collection Center of Pathogenic Microorganisms (CAMS-CCPM-P) until subsequent genome sequencing analysis and following study. Clinical data were collected from the hospital’s computerized medical and biological records concurrently with the collection of the isolates.

### Whole Genome Sequencing and Phylogenetic Analysis

Genomic DNA was extracted using TIANamp Bacteria DNA Kit (TIANGEN) and sequenced on PacBio Sequel platform and Illumina NovaSeq PE150 at Beijing Novogene Bioinformatics Technology Co., Ltd (Beijing, China). The sequencing reads were assembled using SMRT Link v5.0.1 software. Genes were predicted and annotated using GeneMarkS v4.17 (http://topaz.gatech.edu/) and the RAST service (https://rast.nmpdr.org/). MLST analysis was performed using the Multi-Locus Sequence Typing (MLST) v2.0 online service (https://cge.cbs.dtu.dk/services/MLST/) based on whole genome sequencing data. The MLST scheme for *S. aureus* includes seven housekeeping genes (*arcC*, *aroE*, *glpF*, *gmk*, *pta*, *tpi*, and *yqiL*), as described (42). To identify accumulation of variations during infection, we mapped the corresponding long-reads from all 3 strains to the reference genome *S. aureus* CUBIST-27 (phylogenetically closest strain), and then screened for different structural variations (SVs), single nucleotide variants (SNVs), and insertions/deletions (indels) between the strains compared to the reference strain. Detailed variations are listed in Supplementary Table S1. For phylogenetic analyses, an average percentage (AP) distance metric and Neighbour-Joining (NJ) phylogenetic tree was constructed using CLC workbench 21 software as instructed. Detailed accession numbers of genomes for phylogenetic tree construction are listed in Supplementary Table S2.

### Protein Expression and Purification

The gene segment encoding GdpP-C (GdpP313-656) was amplified using ATCC25923 genomic DNA as a template and cloned into the *NdeI* and *XhoI* restriction sites of the pET28a (+) vector. Primer sequences for gene amplification are provided in Supplementary Table S3. Overnight culture was diluted 1:100 into 1 liter LB broth containing 50 μg/mL ampicillin and incubated at 37°C with shaking (220 rpm) until the culture reached an optical density of 0.6-0.8. Expression of GdpP-C-His was induced with 0.2 mM isopropyl β-D-1-thiogalactopyranoside (IPTG) followed by overnight incubation at 20°C with shaking. Bacteria were harvested by centrifugation and resuspended in phosphate-buffered saline (PBS). After sonication, the supernatant was loaded onto a HisTrap™ HP Columns (GE Healthcare) equilibrated with Ni-IDA binding buffer. The column was washed with binding buffer, and then GdpP-C was eluted with Ni-IDA elution buffer (20 mM Tris-HCl, 250 mM imidazole, 0.15 M NaCl, pH 8.0). Finally, the GdpP-C solution was transferred to a dialysis bag and dialyzed overnight in PBS buffer. Protein purity was assessed by SDS-PAGE.

### Western Blot Analysis

Protein expression analysis was performed following a previously reported method (43). *S. aureus* cultures were grown in TSB for 24 hours, harvested by centrifugation (4,500 rpm, 10 min), and lysed with lysostaphin and lysozyme. Protein lysates were dissolved in 5× loading buffer (Beyotime), boiled for 5 min, and separated (10 μg/lane) on 10% precast PAGE gels (Yeasen) at 130 V for 1 hour. Proteins were transferred onto 0.45 μm PVDF membrane (Millipore) using a semi-dry transfer system. Membranes were blocked with 5% non-fat milk (Sangon Biotech) in TBST for 1.5 hours at room temperature. They were then incubated overnight at 4°C with the following primary antibodies: mouse polyclonal anti-α-hemolysin (Hla) (ab190467, Abcam, 1:1,000), rabbit polyclonal anti-protein A (SpA) (bs-0362R, Bioss, 1:1,000), and rabbit anti-GAPDH (5174S, Cell Signaling, 1:1,000). After washing with TBST, membranes were incubated with HRP-conjugated goat anti-rabbit IgG (ZB-2301, ZSGB-BIO) or goat anti-mouse IgG (ZB-2305, ZSGB-BIO) for 1 hour at room temperature. Signals were detected using Omni-ECL chemiluminescence detection reagent (SQ203L, Enzyme) and visualized with an imaging system. Band intensities were quantified using ImageJ software and normalized to β-actin levels. All experiments were performed in three independent biological replicates. One of these replicates is shown in the Result, while the remaining two are presented in Supplementary Figure S1.

### Isothermal Titration Calorimetry (ITC) Assay

The calorimetry assay was performed using the MicroCal PEAQ-ITC (Malvern). Prior to titrations, GdpP-C and pentoxifylline (PTX) were prepared in 50 mM Tris-HCl buffer, respectively. Nineteen injections of 1 mM PTX were titrated into 10 μM GdpP-C at 37°C, with 50 mM Tris-HCl buffer serving as the negative control.

### ATP Quantification Assay

Intracellular ATP levels were measured using a colorimetric ATP assay kit (E-BC-K774-M, Elabscience) according to the manufacturer’s instructions. ATP extraction from bacterial cells was performed following the protocol for *S. aureus*. Briefly, bacterial cultures were harvested by centrifugation at 4°C, and the cell pellets were washed twice with ice-cold PBS. Cells were then lysed using a lysis buffer containing 0.1 M Tris-HCl (pH 7.8), 1% Triton X-100, and 2 mM EDTA with 20 ng/mL lysostaphin. After incubation for 1 h, cell lysis was completed by ultrasonic crushing. The supernatants were collected for ATP determination following centrifugation at 12,000 × g, 4°C for 10 minutes. The ATP content was quantified by measuring absorbance at 460 nm using a microplate reader. ATP concentrations were determined using a standard curve generated from the ATP standards included in the assay kit. Results were normalized to cell pellet wet weight and expressed as nmol ATP per mg wet weight. All measurements were performed in triplicate.

### Quantification of c-di-AMP Levels

Intracellular c-di-AMP concentrations were determined using an ELISA kit (LM-c-di-AMP, Lmai Bio). Sample preparation was performed following a described extraction method with slight modifications. Briefly, bacterial cultures were harvested by centrifugation at 4°C, and the cell pellets were washed twice with ice-cold PBS. Cells were then resuspended in 0.75 mL of 50 mM Tris pH 8 buffer supplemented with 20 ng/mL lysostaphin and incubated for 1 h before completing cell lysis by ultrasonic crushing. Following centrifugation for 5 min at 17,000 × g, the supernatant from the cell lysates were transferred to new tubes. The samples were then heated to 95°C for 10 min, centrifuged for 5 min at 17,000 × g, and the supernatant containing c-di-AMP was transferred to a new tube for analysis using the ELISA kit. Standard curves were generated using the provided c-di-AMP standards, and the measured values were normalized to bacterial wet weight. All measurements were performed in triplicate, and the results are presented as picomoles of c-di-AMP per milligram wet weight.

### GdpP Activity Assay and Pentoxifylline Inhibition Analysis

GdpP enzyme activity was determined using the phosphomolybdate blue colorimetric method in 96-well plates. The reaction mixture (100 μL) contained 50 mM Tris-HCl (pH 7.5), 100 mM NaCl, 5 mM MnCl₂, 0.5 mM DTT, and varying concentrations of c-di-AMP substrate (30-200 μM). Reactions were initiated by adding 10 nM purified GdpP enzyme, incubated at 37°C for 2 minutes, and terminated with 25 μL of 0.5 M EDTA. The released inorganic phosphate was quantified by adding 100 μL of 0.6% ammonium molybdate (in 0.5 M H₂SO₄) and 50 μL of 10% ascorbic acid solution. After 20 minutes of color development at room temperature, absorbance was measured at 700 nm using a microplate reader. Absorbance values were converted to Pi concentrations using a 0-50 μM potassium phosphate standard curve. For pentoxifylline inhibition studies, the enzyme was pre-incubated with 10 μM pentoxifylline for 5 minutes before initiating the reaction. All experiments were performed in triplicate, with three biological and three technical replicates. Enzyme kinetic parameters were calculated by non-linear regression fitting to the Michaelis-Menten equation, and the inhibition type was verified through Lineweaver-Burk double-reciprocal plots. Data are presented as mean ± SD and analyzed using GraphPad Prism 9.0 software.

### MIC Determination

The antimicrobial susceptibilities of the *S. aureus* isolates against trimethoprim-sulfamethoxazole (TMP-SMX) were determined by broth microdilution method, and the interpretative criteria were from Clinical and Laboratory Standards Institute (CLSI) document M100-30th. *S. aureus* ATCC 25923 was used as the quality control strain.

### RNA Extraction and RT-qPCR Gene Expression Analysis

Total RNA of different *S. aureus* strains was extracted from mid-logarithmic phase cultures using RNAprep Pure Cell/Bacteria Kit (TIANGEN, DP430, China). Bacterial cells were collected by centrifugation and processed according to the manufacturer’s instructions. RNA integrity was assessed and the concentration of purified RNA was measured. cDNA was synthesized using FastKing RT Kit (TIANGEN, KR116). RT-qPCR was performed using PowerUp SYBR Green Master Mix (Applied Biosystems, A25742, USA) via a Applied Biosystems™ 7500 Real-Time PCR system (Applied Biosystems, 4351104, USA). The *rpoB* gene was used as an endogenous reference for normalization. Primer sequences for genes of interest and the reference gene *rpoB* are provided in Supplementary Table S3.

### *In vitro* Competition Assay

*In vitro* competition experiments were performed following a previously reported method (44) between strain pairs with different TMP-SMX susceptibility profiles (L1: susceptible; L2 and L3: intermediate). Exponentially growing cells were adjusted to 0.5 McFarland using 0.85% sterile saline, and 10 µL aliquots of each competitor were mixed at a 1:1 (v:v) ratio, followed by 500-fold dilution in 10 mL fresh MH broth. The bacterial mixtures (∼10^5 CFU/mL) were incubated in MH broth at 37°C, 180 rpm for 24 h. Bacterial colonies were enumerated by plating properly diluted cultures on MH agar plates without TMP-SMX for total cell count, and MH agar plates containing 5 μg/mL trimethoprim (TMP component) to selectively quantify the intermediate resistant strains (L2 or L3). This concentration effectively inhibited the growth of the susceptible strain (L1) while allowing growth of strains with intermediate resistance. The relative fitness (RF) was calculated as RF = (lg *S1t* − lg *S1t0*) / (lg *S2t* − lg *S2t0*), where *S1* and *S2* represent CFU densities of the competing strains, and *t* is the measurement time in days. Direct competition experiments were conducted between L1 and L3, and between L2 and L1, while the relative fitness between L2 and L3 was derived indirectly from these results.

### Time-kill Assays

Time-kill assays were performed with clinical *S. aureus* isolates (L1, L2, L3) and reference strain ATCC25923. Assays were conducted in 96-well round bottom plates (Corning Life Sciences, 3799) with 1 mM pentoxifylline in a final volume of 220 μL per well. Overnight cultures were diluted to 10^6 CFU/mL in Mueller-Hinton broth and incubated at 37°C without shaking. Viable cells were counted by plating appropriate dilutions onto LB agar plates at 0, 2-, 4-, 8-, 12-, 24-, 36-, and 48-hours. Results were recorded as lg CFU per milliliter, and all experiments were performed in triplicate on different days.

### Cell Culture and Cell-infection Assays

A549 (human lung epithelial cell) was maintained in DMEM supplemented with 15% FBS without antibiotics at 37°C and 5% CO_2_. For infection assays, cells were seeded in 24-well plates at a density of 2 × 10^5 cells/well 24 hours prior to infection. Before infection, cells were washed twice with PBS and incubated in antibiotic-free medium.

Bacterial strains (L1, L2, L3, and ATCC25923) were grown to mid-logarithmic phase, washed twice with PBS, and diluted in antibiotic-free DMEM. Cells were infected at a multiplicity of infection (MOI) of 10:1 (bacteria:cells), and plates were centrifuged at 1,000 × g for 5 minutes at room temperature to synchronize. After 1 hour of infection, extracellular bacteria were killed by adding gentamicin (100 μg/mL) for 1 hour. Cells were then washed and maintained in medium containing 10 μg/mL gentamicin to prevent extracellular bacterial growth.

To assess intracellular survival, infected cells were lysed at 0, 4-, 8-, 12-, and 24-hours post-infection with 0.1% Triton X-100 in PBS for 10 minutes. Serial dilutions of lysates were plated on appropriate agar for CFU enumeration. All experiments were performed in triplicate.

### Staphyloxanthin Quantitation Assay

Staphyloxanthin was extracted and quantified from *S. aureus* isolates as previously described with modifications (45). Briefly, the tested *S. aureus* strains (L1, L2, L3, and ATCC25923) were grown overnight in LB broth at 37°C, and inoculated to cell density of OD_600_=0.04 in 10 mL LB broth, incubated at 37°C, 220 rpm for 24 h. Staphyloxanthin was extracted with 1 mL methanol from 2 mL cell cultures and then incubated at 55°C with shaking for 30 minutes in the dark, during which the staphyloxanthin was completely dissolved in the methanol. The mixture was centrifuged again at 10,000 rpm for 1 minute to remove the cell pellet. 200 µL of standardized samples were inoculated into each well and repeated three times, and at least three biological replicates per strain were tested. The absorbance at 465nm of the resulting solution was measured to determine relative staphyloxanthin concentration.

### Biofilm Assay

Bacterial strains (L1, L2, L3, and ATCC25923) biofilm assay was performed as described previously with modifications(46). Exponentially growing bacteria were adjusted to 0.5 McFarland using 0.85% sterile saline, followed by 100-fold dilution in fresh Brain Heart Infusion (BHI, BD237500) with 0.5% glucose. Afterwards, the bacterial suspensions (∼10^6 CFU/mL) were allocated into 96-well plates (200 μL/well) and incubated at 37°C for 20 h. After washed twice with 1× PBS (pH 7.4) to remove unattached cells and media components, the biofilm in each well was fixed in the oven at 60°C for 50 min, stained with 200 µL 0.1% crystal violet for 30 min, and washed three times with PBS again and air-dried. The biofilm was then dissolved in 200 µL 95% ethanol for absorbance reading at 590 nm. Every sample was inoculated in sextuplicate, and at least three biological replicates per strain were tested.

### Mouse Infection Model

Three- to five-week-old ICR (CD-1) mice were purchased from Beijing HFK Bioscience Co., Ltd (Beijing, China). All animals were housed under controlled humidity (40 - 60%) and temperature (22 ± 2°C) with a 12 h light-dark cycle. The animal study complied with animal husbandry guidelines, with approval from the Laboratory Animal Welfare and Ethics Committee of the Institute of Medicinal Biotechnology, Peking Union Medical College (Approval number: IMB-20240501D901) Mice were stratified by weight and randomly assigned to three experimental groups: 10 for survival curves, 15 for bacterial counts, and 12 for IL-1β determination. Clinical isolates (L1, L2, and L3) were freshly cultured and adjusted to appropriate density in 0.85% NaCl. Infection was established by intravenous injection of 0.4 mL overnight bacterial culture suspension (∼3.3 × 10˄7 CFU/mL for survival rate comparison; ∼3 × 10˄7 CFU/mL for bacterial burden assessment and IL-1β determination). For survival rate comparison, animal mortality was recorded over 14 days, with survival rates calculated and survival curves plotted. For bacterial burden assessment, blood and organs (liver, spleen, lung, kidney, and brain) were collected 36-, 60-, 72-hours post-infection and homogenized in saline. Bacterial loads were determined using methods previously reported(47). For IL-1β determination, blood was collected at 0, 2-, 4-, 24-hours post-infection and allowed to clot at room temperature for 30 minutes. Samples were then centrifuged at 2,000 × g for 15 minutes at 4°C. The serum was carefully collected. Serum IL-1β levels were measured using a mouse IL-1 beta ELISA kit (KE10003, Proteintech) according to the manufacturer’s instructions.

### Statistical Analysis

Statistical analysis was performed using GraphPad Prism 9.5.1. P values were calculated using t-test or two-way ANOVA multiple comparison test to compare the differences between each pair of groups. A p-value lower than 0.05 was considered statistically significant.

## Data availability

All data supporting the findings of this study are available in the paper and its Supporting Information files. The genome assembly sequence data have been deposited in the NCBI database with accession numbers CP184635 (L1), CP184634 (L2), and CP184633 (L3). Additional data have been deposited in Mendeley Data (DOI: 10.17632/94bjj7ktmz.3). Any further data are available from the corresponding authors upon reasonable request.

## Author Contributions

J.X., J.L., and G.L. designed the research. J.X., L.W., and X.W. performed the experiments. J.X., C.X., Y.M. and Xinyi Y. analyzed the data. J.X., Xuefu Y. wrote the first draft of the manuscript. Xinyi Y., C.L., G.L., and Xuefu Y. revised the manuscript. All authors commented on the manuscript at all stages.

## Acknowledgments

This work was supported by the National Natural Science Foundation of China (82330110, 32141003), the National Key Research and Development Program of China (2024YFC2309300), the CAMS Innovation Fund for Medical Sciences (2021-I2M-1-026, 2021-I2M-1-039), and the National Science and Technology Infrastructure of China (National Pathogen Resource Center-NPRC-32).

## Institutional Review Board Statement

Our study was approved by the Ethics Review Committee of The First Affiliated Hospital of Hebei North University (Approval number: K2024410, approved on Apr. 10, 2024). As this was a retrospective study of clinical electronic records and bio-specimens that did not involve identifiable human material or data, the requirement for patient informed consent was waived by the Ethics Committee.

## Conflicts of Interest

The authors declare no conflicts of interest.

---

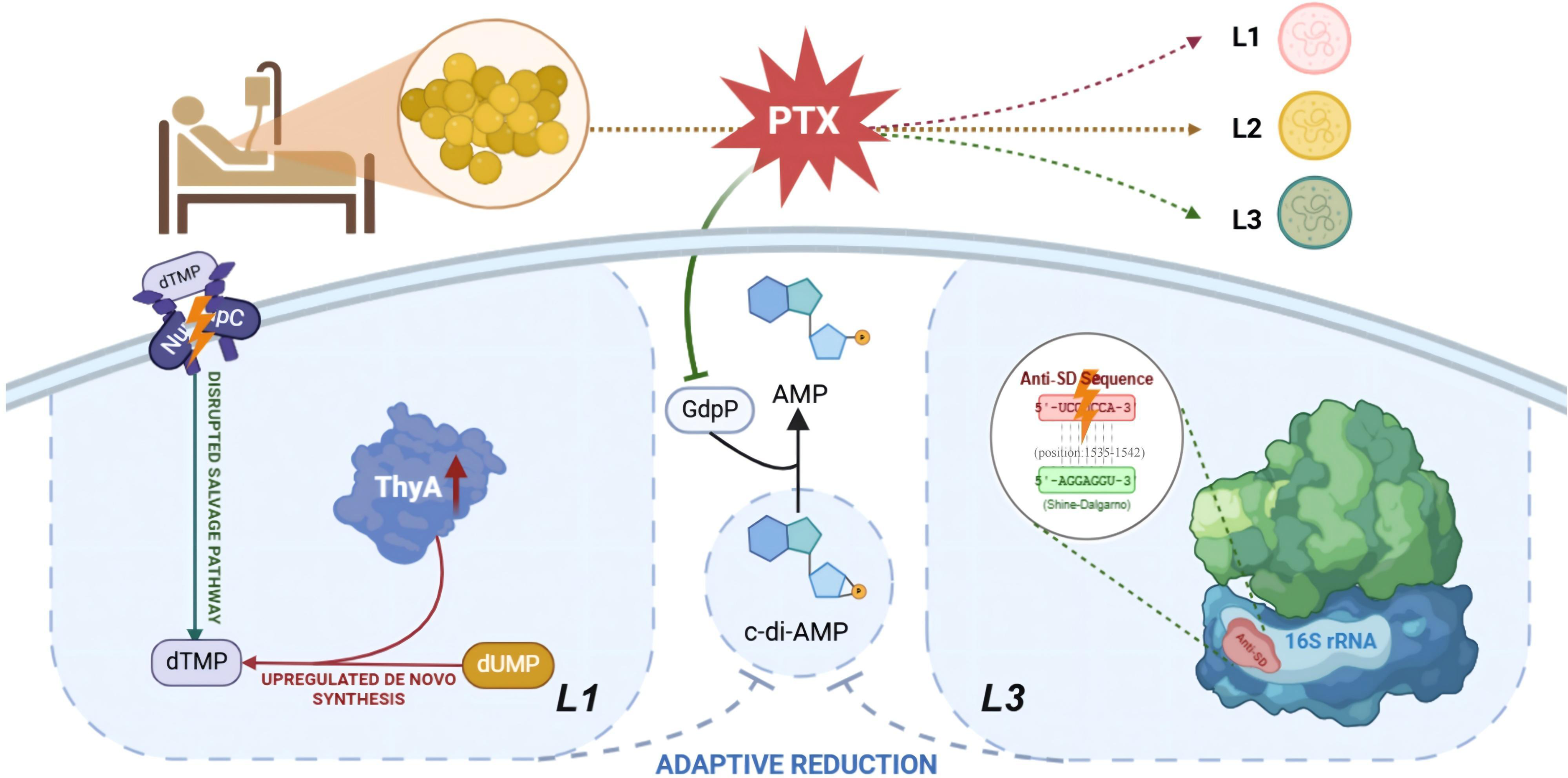

